# Examining the genetic influences of educational attainment and the validity of value-added measures of progress

**DOI:** 10.1101/233635

**Authors:** Tim T Morris, Neil M Davies, Danny Dorling, Rebecca C Richmond, George Davey Smith

## Abstract

In this study, we estimate (i) the SNP heritability of educational attainment at three time points throughout the compulsory educational lifecourse; (ii) the SNP heritability of value-added measures of educational progress built from test data; and (iii) the extent to which value-added measures built from teacher rated ability may be biased due to measurement error. We utilise a genome wide approach using generalized restricted maximum likelihood (GCTA-GREML) to determine the total phenotypic variance in educational attainment and value-added measures that is attributable to common genetic variation across the genome within a sample of unrelated individuals from a UK birth cohort, the Avon Longitudinal Study of Parents and Children. Our findings suggest that the heritability of educational attainment measured using point score test data increases with age from 47% at age 11 to 61% at age 16. We also find that genetic variation does not contribute towards value-added measures created only from educational attainment point score data, but it does contribute a small amount to measures that additionally control for background characteristics (up to 20.09% [95%CI: 6.06 to 35.71] from age 11 to 14). Finally, our results show that value-added measures built from teacher rated ability have higher heritability than those built from exam scores. Our findings suggest that the heritability of educational attainment increases through childhood and adolescence. Value-added measures based upon fine grain point scores may be less prone to between-individual genomic differences than measures that control for students’ backgrounds, or those built from more subjective measures such as teacher rated ability.

## Introduction

Researchers and policy makers frequently use value added (VA) measures to quantify educational progress irrespective of a child’s ability, by comparing performance at a given stage of education to performance at an earlier stage^1^. Because VA measures use a baseline level of attainment they should control for between-individual time-invariant differences such as inherent ability. This means that they offer a measure of educational progress that is free from confounding by such factors, and therefore do not unfairly assess the final grades of schools with more disadvantaged student intakes^2^. VA measures are therefore – in theory - a useful tool for assessment of educational services. Further data can be used to supplement the raw VA measures and adjust for additional factors that influence education to create contextual value-added (CVA) measures (see Box 1). There has however been debate over which background factors should be adjusted for and the extent to which CVA measures sufficiently adjust for these factors^3^. VA measures are widely used in educational research to provide a measure of teacher and school effectiveness, with research showing that children assigned to high-VA teachers are more likely to enter post-compulsory education and earn higher salaries in later life^4^. They have played an important role in determining the published performance of individual schools in UK school league tables - themselves a highly contentious issue^2^ - and have directly influenced school accountability^5^.

#### Box 1: Difference between raw value-added measures and contextual value-added measures

##### Raw value-added

Raw value-added measures are used to assess the contributions that contextual factors such as teachers and schools make towards a child’s education progress. They compare a child’s test score at a given stage of education to the same child’s test score at a previous stage of education. Because the baseline test score inherently contains a measure of child ability, raw value-added measures are seen to offer a measure of educational progress that is free from confounding by time-invariant factors. This makes them a fairer measure of comparison for the effectiveness of teachers and schools than raw attainment scores, which are confounded by earlier attainment and therefore biased measures of progress.

##### Contextual value-added

Contextual value-added measures are similar to raw value-added measures but they account for a range of additional time-invariant factors. These factors may include variables such as gender, ethnicity, special educational needs, month of birth, and so on. By adjusting for these variables, contextual value-added measures are seen to further control for factors that are beyond the teacher or schools control in addition to child ability.

Because they account for all time-invariant factors between children, VA measures are considered (by many of their proponents) to be a reliable and unbiased way to measure school and teacher performance independent of selection of pupils, background characteristics and innate ability^6^. However, there has been criticism of VA measures and debate over the extent to which they successfully control for time-invariant factors^7,8^. A key factor that VA measures are designed to overcome is genetic difference between individuals; the genetic variants that we inherit are fixed and therefore do not vary over time. This is important because there is a large body of evidence suggesting that genetic variation affects educational attainment when defined using either narrow or broad sense heritability^9–12^. Narrow sense heritability is defined as the proportion of total variance in a trait that is due *only* to additive genetic variance, while broad sense heritability is the total proportion of variance in a trait that is due to total genetic variance, *inclusive* of additive genetic variance, dominance, epistasis (gene-gene interactions) and parental effects. Meta-analyses of twin studies has revealed an overall educational attainment broad sense heritability of 40% (95% CI: 35–45%)^11^, and the largest GWAS to date has demonstrated the polygenicity of attainment^13^. There has been considerable variation in results; using data from the Twins Early Development Study (TEDS)^1^ narrow sense heritability of attainment at age 16 has been estimated at 31%^14^ while broad sense heritability was estimated at 62%^12^.

Recent genetic analyses have suggested that VA measures are indeed influenced by genetic differences between children. An analysis of TEDS estimated VA score broad sense heritability to be 52% (95% CI: 48–57). This finding led Haworth and colleagues (2011) to conclude that VA measures are as strongly influenced by genetic factors as raw attainment scores and therefore that they perform poorly at controlling for time-invariant differences between children. The study by Haworth and colleagues used teacher reported ability instead of the directly assessed attainment scores which are used to inform school league tables and educational policy. The choice of assessment measure is important because teacher rated ability is likely to be a less accurate measure of student achievement than test-assessed point scores, and characterised by systematic error due to confounding by teacher-reported bias on traits such as ethnicity or physical attractiveness^15–17^. Therefore, the previous study’s heritability estimates may have been biased using teacher assessed ability. Further investigation into the ability of value-added measures to control for genomic differences between children is therefore required to assess the suitability of VA measures for informing research and policy.

Molecular genetic data offers an opportunity to examine the narrow sense heritability of educational attainment, and to test the ability of VA measures to control for time-invariant between individual differences (one of their desirable properties). If VA measures are estimated to have non-zero heritability then they are susceptible to the influence of genetics. Throughout this paper we use the term *heritability* to refer to SNP-based heritability, a measure of narrow sense heritability calculated from a given set of SNPs. In this study we (i) estimate the heritability of educational attainment at three time points throughout the compulsory educational lifecourse; (ii) estimate the heritability of raw value-added (VA) and contextual value-added (CVA) measures created from examination assessment data to assess if they control for time-invariant between individual genetic differences; (iii) test if these heritabilities differ from VA measures created from teacher assessed ability.

## Results

Table 1 displays the number of study children who provide information for each of the analyses and descriptive statistics for each of the variables used. Sample size is higher for the raw attainment measures but reflects some data loss in the raw VA and CVA measures due to missing attainment and background factors respectively. The CVA measures have higher standard deviations than the raw VA measures because they are measured using differences between predicted point scores and realised point scores, whereas the raw VA measures use differences in standardised score differences.

**Table 1:** Descriptive statistics for children included in the analyses

|  | n | Mean | SD |
| --- | --- | --- | --- |
| KS2 points | 6132 | 28.04 | 3.85 |
| KS3 points | 4960 | 35.97 | 6.19 |
| KS4 points | 6518 | 39.89 | 9.48 |
| KS 2-3 VA score | 4904 | 0.06 | 0.42 |
| KS 2-4 VA score | 6088 | -0.03 | 0.63 |
| KS 3-4 VA score | 4924 | -0.07 | 0.49 |
| KS 2-3 CVA score | 4600 | 0.10 | 2.52 |
| KS 2-4 CVA score | 6028 | 0.98 | 56.33 |
| KS 3-4 CVA score | 4914 | 1.08 | 45.21 |

SD, standard deviation; VA, raw value-added; CVA, contextual value-added.

### Heritability of traits

The SNP heritabilities of educational attainment measured at each of the Key Stages are presented in Figure 1. Heritability rises with age from 47.3% (95%CI: 35.9 to 58.7) at age 11 to 57.6% (95%CI: 43.9 to 71.3) at age 14 and 61.1% (95%CI: 50.7 to 71.5) at age 16. These heritabilities are higher than would be expected given the results from a meta-analysis of educational attainment suggesting a heritability of 40.0% (95% CI: 35.3–44.7)^11^. This suggests that in the ALSPAC sample genetic variation contributes towards around half of the total variance in educational attainment using exam scores.

**Figure 1:**
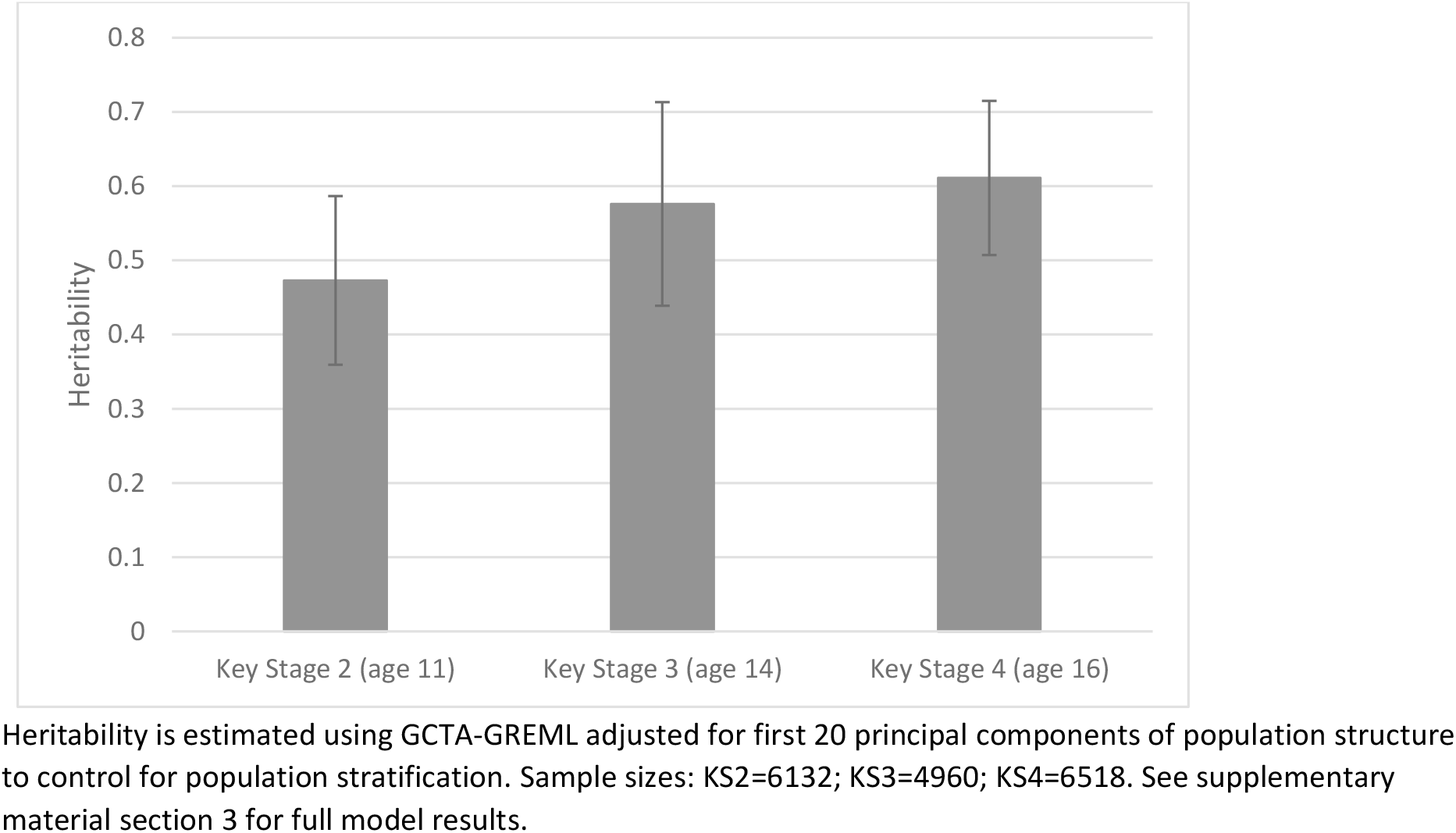
Heritability of Key Stage average point score attainment

### Value-added measures

Figure 2 displays the heritability of VA measures. There is a very small amount of genetic variance in the raw VA measures, far smaller than for the raw attainment scores at each Key Stage. The raw VA measures from ages 11 to 14, 11 to 16, and 14 to 16 show SNP heritabilities of <0.1%^2^ (95%CI: −13.5 to 13.5), 7.9% (95%CI: −3.3 to 19.0), and 6.5% (95%CI: −7.6 to 20.6) respectively, although in each case the standard errors are large in comparison to the estimates resulting in little evidence of heritability in the raw VA measures. The heritability estimates for the CVA measures (Figure 2) are consistently higher than those for the corresponding raw VA measures. There is evidence for a small amount of heritability in the CVA scores for the period of age 11 to 14 (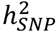: 20.09%; 95%CI: 6.06 to 35.71), and the period of age 11 to 16 (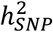: 15.77%; 95%CI: 4.26 to 27.29). Heritability estimates for the period from age 14 to 16 are lower with no strong evidence for a heritable component (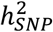: 8.49%; 95%CI: −5.52 to 22.51). These results consistently suggest that relatively little of the variation in raw VA measures – the unadjusted progress that students make from one Key Stage to another – can be attributed to common genetic variation. However, they also suggest that the variation in some CVA measures – those that adjust for background factors – *can* in part be attributed to common genetic variation. To explore this somewhat counterintuitive finding we ran a series of simulations to determine scenarios in which CVA measures may be more genetically biased than raw VA measures (see supplementary material section 1). The simulations demonstrate that CVA measures overstate heritability more than raw VA measures where the baseline input score contains measurement error (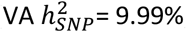; 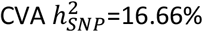 given measurement error of 0.25).

**Figure 2:**
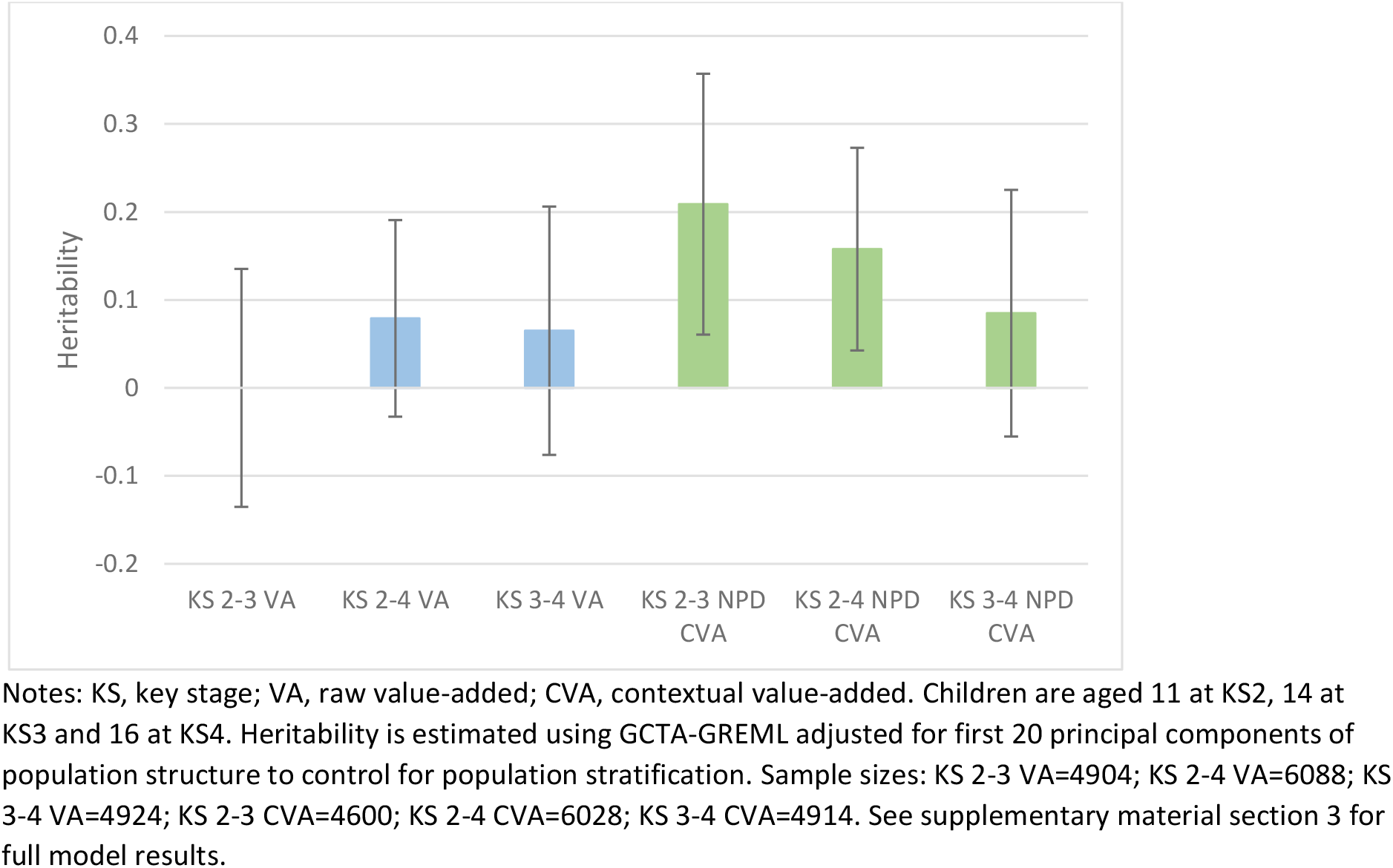
Heritability of Value-added measures

These results contrast with those from a previous study which found that VA measures using teacher assessed ability were around 50% heritable^18^. While teacher assessed ability may be a good proxy for true student ability, it may be confounded by teacher-report bias which will not be present in directly assessed examination scores. If baseline achievement is measured with error due to teacher-reported bias, then it will not fully control for prior achievement. Furthermore, because teacher assessed ability is based on broader scales (fewer bandings) than the rich point scores used by exams assessments, the variability of scores is lower as children cannot be differentiated within the bands. Where the baseline measures are more constrained they will be less effective at controlling for initial differences, and will potentially lead to upward biased estimates of heritability. It is therefore possible that the high heritability observed in the previous study may be due at least in part to a lack of measurement precision or bias on the input variable compared to point scores that are traditionally used in educational research.

We investigated if the discrepancy between our results and those from the previous study may have reflected genuine differences (a sample issue) or differences caused by alternative methods of assessment (a measurement issue). A teacher assessed value-added (TAVA) measure similar to that used in the previous study was created for the period of education from age 11 to age 14^3^. The ages that assessments were made were similar for both samples (ages 11 and 14 in ALSPAC and ages 10 and 12 in TEDS). Figure 3 presents the GCTA heritability results of our teacher assessed value-added (TAVA) measure compared to the raw VA and CVA measures at this age. The results suggest a moderate amount of heritability in the teacher assessed measure of 36.3% (95%CI: 22.8 to 49.8). This exceeds the heritability estimates presented in Figures 1 and 2 and suggests that VA measures using teacher rated ability are likely to reflect both student progress and genetic differences between students.

**Figure 3:**
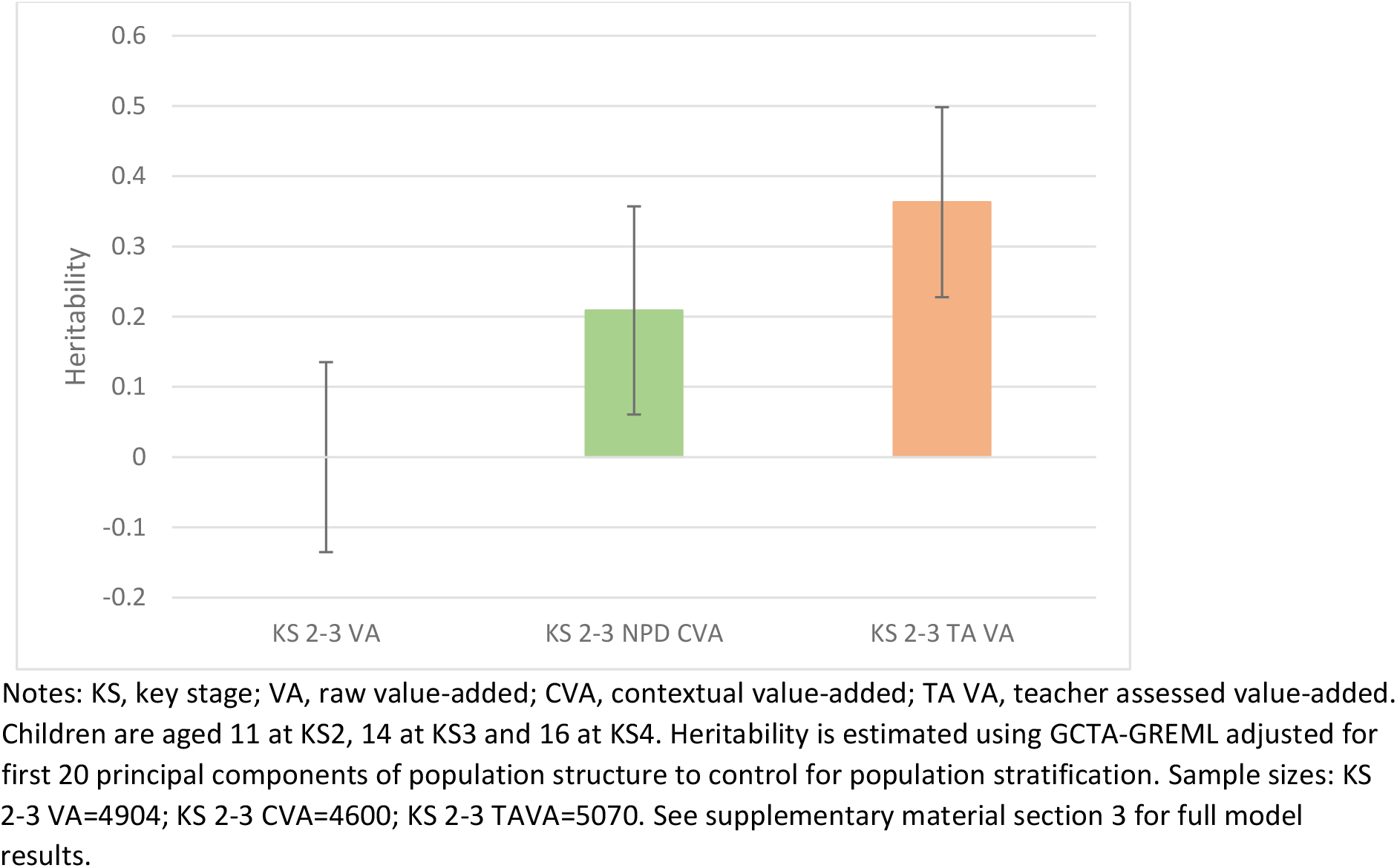
Heritability of KS2-3 Value-added measures

### Polygenic score results

The final stage of the analysis was to investigate whether the educational attainment polygenic score (EAPGS)^13^, that is, the combination of individual genetic variations known to influence educational attainment, predict variance in educational attainment and VA measures. The results (Figure 4) show that the EAPGS accounts for a small but detectable proportion of variance in educational attainment at each Key Stage, varying from 0.35% to 0.58%. The EAPGS accounts for negligible variation in the raw VA measures (≤0.01%), providing further evidence that these successfully account for time-invariant differences between children and are not influenced by genetic factors. Consistent with the GCTA results the EAPGS predicts a small but detectable amount of variation in the CVA measures (0.07% to 0.21%), though this is far smaller than the raw attainment scores. Again, this suggests that these adjusted CVA measures are more strongly associated with genetic differences than raw VA measures. The EAPGS explains a greater amount of variation in the TAVA measure (0.25%) than the raw VA (<0.01%) and CVA (0.07%) measures for this period, closer to the heritability of the age 11 and 14 point scores themselves. Given that the EAPGS only accounts for 74 variants this provides quite remarkable further evidence that value-added measures based upon teacher rated ability are likely to be considerably biased by genetic variation associated with educational attainment.

**Figure 4:**
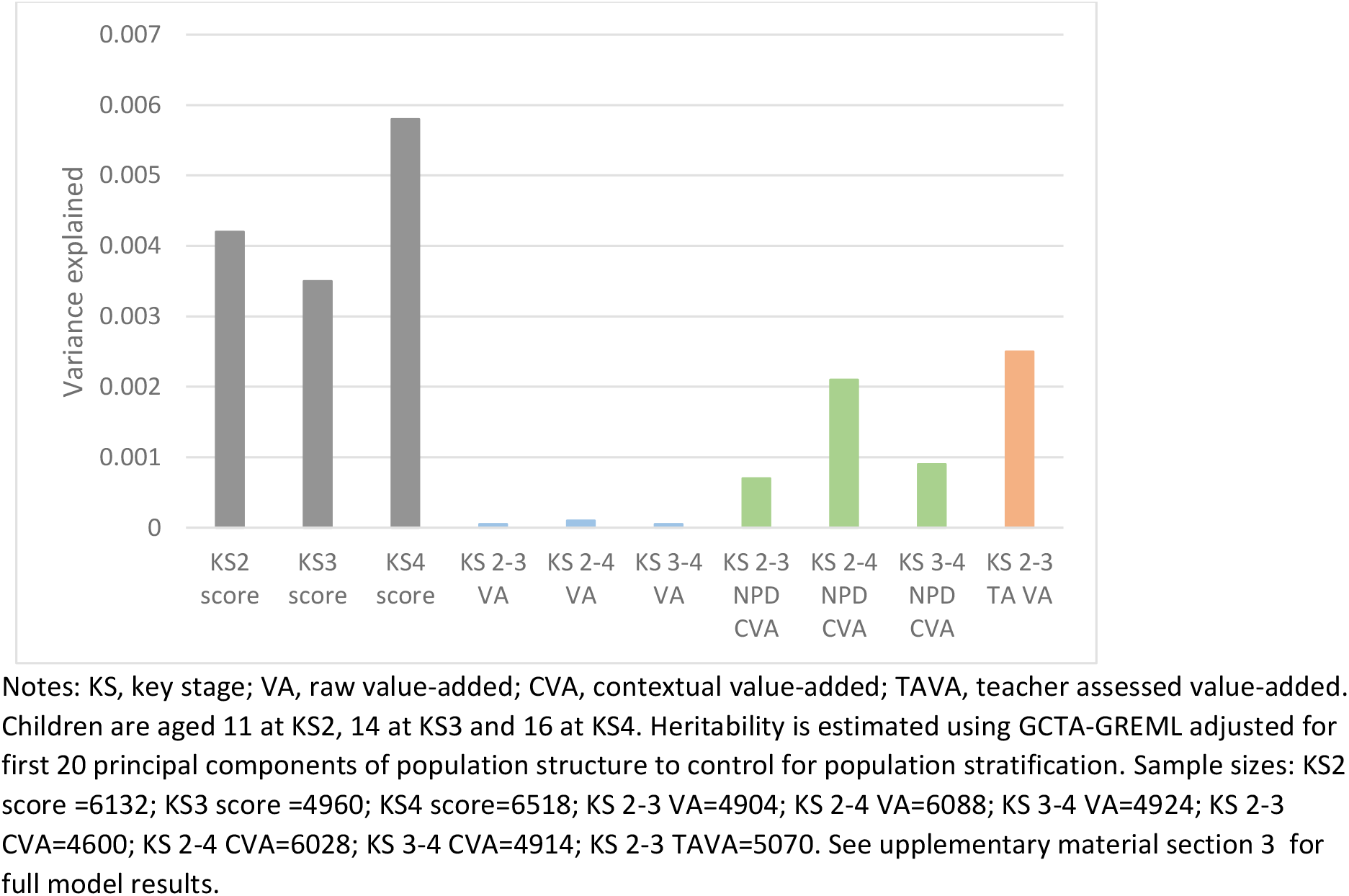
Proportions of variance in outcomes attributed to the EAPGS

## Discussion

### Attainment throughout the educational lifecourse

Our results provide evidence of increasing heritability of educational attainment with age in the same cohort, conforming to previous findings. Our genome wide heritability estimates taking account of all common genetic variants suggest that the heritability of educational attainment increases from 47% at age 11 to 61% at age 16. This increase suggests that environmental factors account for fewer differences between scores at age 16 than earlier ages. This is important because the age 16 exams play a large role in setting a child’s chances in further education and the labour market. The differences across ages are small and could be due to various factors. Parents, teachers and others may work harder at later ages to remove environmental influences, making them more homogenous earlier ages. Attitudinal behaviours may also play a part because older children have more freedom to choose their own educational effort with less parental influence. It is also important to note that in some subjects such as Maths, children were entered into different tiers after the age 14 exams, which placed ceiling and floor caps on the scores that they could attain in the age 16 examinations^4^.

Our estimated heritability is higher than the heritability of 40.0% (95% CI: 35.3 to 44.7) estimated using meta-analyses^11^, and similar to the broad sense heritability of 62% (95% CI: 58% to 67%) in the TEDS cohort for age 16 grades^12^. Our estimate also exceeds than the narrow sense heritability of 31% (95% CI: 7.48% to 54.52%) for age 16 grades estimated using GCTA-GREML in TEDS^14^. It is possible that the heritability of point scores may be higher than that of years of education (as used in the meta-analysis) or grades (as used in TEDS), but it must be noted that this may also reflect an estimation issue between broad and narrow sense heritability. The difference may also reflect the fact that the ALSPAC cohort experience more homogenous environments than other cohorts used in the meta-analysis; all the children in ALSPAC were born within three years in the same geographical location and they mostly experienced the same school system. This may reduce the amount of environmental variation, which will increase the proportion of variation in the phenotype that can be attributed to common genetic variation.

### Value-added measures

We found that very little of the variation in raw value-added measures built from point score data could be attributed to common genetic variation. Surprisingly, we found that these raw VA measures outperformed CVA measures which additionally control for background factors. This suggests that raw VA measures may be less prone to between-individual genomic differences than CVA measures and therefore offer a more reliable measure for the value added by schools or teachers. Our simulations suggest that the reduced performance of CVA measures may be due to measurement error in the baseline scores which results in inflated genetic influences. Because KS2 and KS3 point scores are determined from a smaller range of subjects than KS4 point scores, they are likely to provide baseline measures of overall academic ability which contain greater measurement error. This will therefore result in CVA measures being more biased by genetic influences than raw VA measures. Issues with controlling for background factors in measures of educational progress have long been noted, with some arguing that adjusting for factors may increase bias where they are crude proxies, broad input variables, or related to parental choice^5^^3^. These simulations may partly explain why VA measures built from teacher reported ability further overestimate heritability as they are likely to contain greater measurement error than VA measures built from point scores. Our findings therefore offer some support for using VA measures in educational research, which are largely independent of genetic background, though caution should be taken when considering the construction of the VA measure.

The VA measures created from teacher rated ability for the same children show considerably higher heritability, of around 36%, suggesting that teacher rated ability is more prone to between-individual genomic differences than official test point scores data. Several factors may account for this higher heritability estimate. For example, there is evidence that teacher rated ability may be influenced by heritable factors such as attractiveness which could lead to confounding bias^19,20^. Furthermore, teacher rated ability is likely to be a less precise measure of ability than end of key stage point scores due to its reduced richness as a measure of attainment and therefore reduced imposed variability. If teacher rated prior attainment is measured with bias or low precision, then VA measures based on this will not fully account for baseline differences between children. As our simulations demonstrate, this may upwardly bias heritability estimates. In contrast, Key Stage exams are likely to measure attainment with less error and therefore provide VA measures that suffer from lower bias.

Our teacher rated VA estimate contrasts with that from the only previous study examining genetic influences on VA measures which estimated heritability in TEDS at 50%^18^.It possible this discrepancy is because the two cohorts represent different samples. However, the two cohorts are drawn from the same country and the age of participants differs only by a maximum of 6 years. It is therefore unlikely that this leads to the observed differences (assuming teachers of the geographically concentrated ALSPAC study were as accurate at determining student ability as the general population of teachers). Second, it is possible that the teacher reported ability collected by TEDS may have been less accurate or subject to greater bias than those linked from the UK National Pupil Database (NPD) to the ALSPAC cohort. It is likely that a child’s level would have been decided with more care in the official (and contractually required) National Curriculum reports used by ALSPAC than the optional survey used by TEDS. Third, the ALSPAC analytical sample is likely to be more genotypically homogenous than the TEDS sample because it is geographically concentrated rather than national. Fourth, it is also possible that the discrepancy between the TEDS and ALSPAC samples is due to the age at which teacher reported ability was measured. ALSPAC measures were taken at ages 11 and 14 (the end of Key Stages 2 and 3) while the TEDS measures were taken at ages 10 and 12. Because ALSPAC has no data on teacher reported ability at age 16 (Key Stage 4) we were unable to test for variation in teacher reported value-added measures across the educational lifecourse in the same way as raw and contextual value-added measures. It is therefore possible that such variation may be in part driving the observed differences between the two cohorts. It is also possible that this discrepancy in findings is at least in part due to differences between GCTA and twin models, however the discrepancy is likely too large to be fully accounted for by these model differences. Ultimately, further work replicating the GCTA approach is required on other datasets such as TEDS to resolve these differences and determine if they are due to differences in modelling approach or differences in the data measures used.

### Polygenic scores

We found that the EAPGS consisting of 74 SNPs explained between 0.35% and 0.58% of the variation in educational attainment depending on the stage of education. The EAPGS explains far less variation in exam scores than the GCTA estimates because it uses a set of only 74 genetic variants that associate with education at genome-wide levels of significance p<5e^−8^, a tiny proportion of the genome wide variants used by GCTA, and therefore has less explanatory power. This is demonstrated by two recent studies using TEDS data; a genome wide score using 108,737 SNPs explained 9% of the variance in age 16 grades^21^ whereas a score using only 5,733 SNPS explained only 1.5% of the variance in the same trait^14^. The polygenic score results corroborated with the GCTA results for the value-added measures.

The EAPGS explained negligible variance in the raw VA measures (≤0.01%), suggesting that the 74 variants in the score did not associate with progress measured using only attainment. However, the EAPGS explained a greater amount of variance in the CVA measures (0.07% to 0.21%) and the teacher rated VA measure (0.25%), highlighting that these measures of progress are prone to between-individual genomic differences in educational attainment. These results further demonstrate that value-added measures which adjust for background variables or those built from teacher rated measures of ability may be confounded by genetics, and therefore unsuitable for fairly assessing teachers or schools.

### Limitations

These findings provide only estimates of the variance in educational attainment and value-added measures across the lifecourse, and do not imply that common genetic variants determine the educational attainment of an individual. The major limitation with this work relates to the potential of GCTA to overestimate heritability, which can occur where model assumptions, particularly that of even linkage disequilibrium between SNPs, are violated^22^. Furthermore heritability estimates from GCTA are sensitive to the sampling of participants, the accuracy of phenotypic measurement, and the structure of the genetic relatedness matrix underlying the data^23^. These limitations however have been strongly refuted by the authors of GCTA^24,25^. Nevertheless, our estimates are comparable to those previously conducted using twin designs despite SNP heritability typically being lower than that derived from twin studies. It is therefore possible that our high heritability estimates may suffer from over inflation. One further issue that may lead to overestimation of heritability estimates is that of dynastic effects, where the parental phenotype/genotype directly affects the offspring’s outcomes. It has recently been demonstrated that this can upwardly bias GCTA heritability estimates of educational attainment due to the methods inability to distinguish between direct and indirect genetic effects^26^. There may also be unobserved differences between individuals (residual population structure) biasing our results; we attempted to account for this by using the first twenty principle components of population structure however, we cannot be certain that these will correct for all differences. This limitation could be overcome with genotypic data on mother-father-offspring trios and future studies should exploit the growing availability of data to investigate this hypothesis. Given that data on teacher reported ability was only available for ages 11 and 14 we were unable to examine if the bias in value-added measures based upon teacher reported ability was consistent between the ages of 11 to 16 and 14 to 16. Future studies with teacher reported ability at multiple timepoints should examine any such variation by age. As with all measures based upon educational attainment, random measurement error at the individual level will exist within the data (such as a child suffering illness at the time of examination). However, these random changes will likely provide only a minimal amount of bias at the aggregate level given our sample size. Conversely, teacher rated measures could be susceptible to longer term factors that may impact a child’s educational performance such as parental illness. Finally, it is inevitable that our measure of value-added will still contain measurement error. Random measurement error will not be related to genetic (or other) underlying factors and could therefore bias heritability estimates towards the null. However, it is unlikely that our raw VA measure will suffer from heavy bias because the NPD Key Stage examinations represent the most accurate objective assessment of a child’s educational ability.

### Concluding remarks

In conclusion, our results demonstrate that the heritability of educational attainment – measured using national standardised fine graded point scores - is relatively consistent throughout the educational lifecourse, ranging from 47% to 61%. Our results also suggest that raw value-added measures successfully control for genomic differences between children and outperform contextual value-added measures, and that both measures built from fine grade point scores outperform VA measures based upon teacher reported ability. Value-added measures that underpin school league tables and inform school and teacher accountability may therefore be confounded by genetics and provide unfair assessments of teachers or schools.

## Methods and materials

### Study sample

Participants were children from the Avon Longitudinal Study of Parents and Children (ALSPAC). Pregnant women were eligible to enrol if they had an expected date of delivery between April 1991 and December 1992 and were resident in the (former) Avon Health Authority area in South West England (for full details of the cohort profile and study design see Boyd et al (2013) and Fraser et al (2013))^6^. The ALSPAC cohort is largely representative of the UK population when compared with 1991 Census data; however there is under representation of some ethnic minorities, single parent families, and those living in rented accommodation. Ethical approval for the study was obtained from the ALSPAC Ethics and Law Committee and the Local Research Ethics Committees. From the core sample of 14,775 children 14,115 have data on at least one measure of educational attainment. From these children genetic data was available for 7,988 after quality control and removal of related individuals. We utilise the largest available samples in each of our analyses to increase precision of estimates, regardless of whether a child contributed data to the other analyses.

### Education variables

#### Educational attainment

Our measures of educational attainment are average fine graded point scores at each of the major stages of education in the UK; Key Stage 2 assessed at age 11; Key Stage 3 assessed at age 14; and Key Stage 4 assessed at age 16. Point scores were used to obtain a richer measure of a child’s attainment than level bandings, with the distributions of the raw scores presented in the supplementary material section 2. Scores for the ALSPAC cohort were obtained from the UK National Pupil Database through data linkage which represents the most accurate record of individual educational attainment available in the UK. We extracted all scores from the Key Stage 4 database as this includes attainment at earlier Key Stages and provides a larger sample size than the earlier databases.

#### Educational attainment polygenic score

From these genome-wide data, an educational attainment polygenic score was generated based upon the 74 SNPs identified in the largest GWAS of education to date^13^. The dose of the effect allele at each locus was weighted by the effect size of the variant in the replication cohort of the meta-analysis, the UK Biobank^7^, and these doses were summed using the allelic scoring function in PLINK (version 1.9)^29^.

#### Genetic data

ALSPAC children were genotyped using the Illumina HumanHap550 quad chip genotyping platforms by 23andme subcontracting the Wellcome Trust Sanger Institute, Cambridge, UK and the Laboratory Corporation of America, Burlington, NC, US. The resulting raw genome-wide data were subjected to standard quality control methods. Individuals were excluded on the basis of gender mismatches; minimal or excessive heterozygosity; disproportionate levels of individual missingness (>3%) and insufficient sample replication (IBD < 0.8). Population stratification was assessed by multidimensional scaling analysis and compared with Hapmap II (release 22) European descent (CEU), Han Chinese, Japanese and Yoruba reference populations; all individuals with non-European ancestry were removed. SNPs with a minor allele frequency of < 1%, a call rate of < 95% or evidence for violations of Hardy-Weinberg equilibrium (P < 5E-7) were removed. Cryptic relatedness was measured as proportion of identity by descent (IBD > 0.1). Related subjects that passed all other quality control thresholds were retained during subsequent phasing and imputation. 9,115 subjects and 500,527 SNPs passed these quality control filters.

Children’s genotypes were jointly phased and imputed with the genotypes of the ALSPAC mothers (Illumina human660W quad (mothers)), combining 477,482 SNP genotypes which were in common between the samples. SNPs with genotype missingness above 1% were removed due to poor quality (11,396 SNPs removed) and a further 321 subjects due to potential ID mismatches. This resulted in a dataset of 17,842 subjects containing 6,305 duos and 465,740 SNPs (112 were removed during liftover and 234 were out of HWE after combination). Haplotypes were estimated using ShapeIT (v2.r644) which utilises relatedness during phasing. We obtained a phased version of the 1000 genomes reference panel (Phase 1, Version 3) from the Impute2 reference data repository (phased using ShapeIt v2.r644, haplotype release date Dec 2013). Imputation of the target data was performed using Impute V2.2.2 against the 1000 genomes reference panel (Phase 1, Version 3) (all polymorphic SNPs excluding singletons), using all 2186 reference haplotypes (including non-Europeans). This gave 8,237 eligible children with available genotype data after exclusion of related subjects using cryptic relatedness measures.

#### Value-added measures

We use two sets of value-added (VA) measures in our analyses. First, we calculated a raw VA score as the difference between standardised point scores at different Key Stages. This VA measure can be considered the child’s cohort specific VA score as it is based upon the rank ordering of the child in the cohort at each occasion. Second, contextual value-added (CVA) measures were extracted from the National Pupil Database (NPD) database linked to ALSPAC participants. The CVA measures - using the example of a CVA score between ages 11 and 14 - are calculated as the difference between a child’s given exam score (age 14) and the score that would be predicted from that child’s previous Key Stage exam score (age 11). The models used to calculate CVA measures are estimated within a multilevel framework whereby students are nested within schools and the intercept is permitted to vary across schools. The CVA models also account for gender; special educational needs; eligibility for free school meals (a proxy for low income); first language; school mobility; ethnicity; month of birth; an indicator of whether a child has been in care; and residential area level deprivation. We present results for both the CVA and VA measures throughout our analyses.

We also use a value-added measure based on teacher-assessed ability of children. Teachers are required to grade their students at multiple time points in English, Mathematics and Science on a scale of 1 to 8. These grades reflect the level at which a teacher deems a student to be working, with higher levels reflecting students working at a more advanced stage. Because the levels run throughout a child’s educational career they can be compared at different time points to assess progress. The TAVA measure we use was calculated as the difference between the mean level of English, Maths and Science at ages 11 and 14, thus representing progress during these years. Teacher-reported ability is not available in the NPD at age 16, meaning that our teacher assessed value-added (TAVA) measure only covers the one educational period.

### Statistical analysis

To estimate the proportion of variation in educational attainment and VA measures that can be attributed to common genetic variation (the SNP heritability) we use a series of univariate analyses as follows:

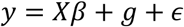

where *y* is the heritability of a phenotype, *X* is a series of covariates, *g* is a normally distributed random effect with variance 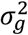, and *ϵ* is residual error with variance 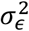. Heritability is defined as the proportion of total phenotypic variance (genetic variance plus residual variance) attributed to genetic variation:

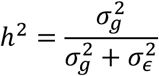

All analyses were conducted using whole-genome restricted maximum likelihood (GREML) in the software package GCTA^8^, with results adjusted for the first 20 principal components to control for population stratification^9^. The GCTA software uses measured SNP level variation to estimate the genetic similarity between every two unrelated individuals and compares this to the phenotypic similarity between them. If genetically similar pairs are more phenotypically similar than genetically dissimilar pairs then heritability estimates of the phenotypic trait in question will be higher. This analysis is restricted to unrelated participants (less related than 2^nd^ cousins), as indicated by the ALSPAC Genetic Relatedness Matrices. All continuous variables were inverse-normally-rank-transformed to have a normal distribution.

## Funding

The Medical Research Council (MRC) and the University of Bristol fund the MRC Integrative Epidemiology Unit [MC_UU_12013/1, MC_UU_12013/9]. NMD is supported by the ESRC via a Future Research Leaders Fellowship [ES/N000757/1].

## Acknowledgements

We are extremely grateful to all the families who took part in this study, the midwives for their help in recruiting them, and the whole ALSPAC team, which includes interviewers, computer and laboratory technicians, clerical workers, research scientists, volunteers, managers, receptionists and nurses. Ethical approval for the study was obtained from the ALSPAC Ethics and Law Committee and the Local Research Ethics Committees. The UK Medical Research Council and the Wellcome Trust (Grant ref: 102215/2/13/2) and the University of Bristol provide core support for ALSPAC.

## Footnotes

1 See https://www.teds.ac.uk/

2 The estimate of heritability for the KS2-3 raw VA score is constrained to zero due because the predicted heritability is −0.04, which is outside of the bounds of zero to one.

3 Teacher rated ability was only available at age 11 (KS2) and age 14 (KS3)

4 For a brief overview of tiering in age 16 exams see^30^

5 For example, if high SEP parents respond to poor baseline achievement by increasing family inputs, a correlation between family socioeconomic status and educational progress would be induced^3^.

6 The study website contains details of all the data that is available through a fully searchable data dictionary (ALSPAC data dictionary available at http://www.bris.ac.uk/alspac/researchers/data-access/data-dictionary/).

7 See http://www.ukbiobank.ac.uk/

8 See http://cnsgenomics.com/software/gcta/

9 Population stratification refers to the way in which different subpopulations may have systematic differences in allele frequencies due to ancestral differences. Principal components of inferred population structure are included as covariates in analyses to account for these population specific variations in allele distributions.

## Notes

The authors declare no conflict of interest. This publication is the work of the authors and Tim Morris will serve as guarantor for the contents of this paper.

